# A Multiscale Framework for Deconstructing the Ecosystem Physical Template of High-Altitudes Lakes

**DOI:** 10.1101/034405

**Authors:** Dragos G. Zaharescu, Peter S. Hooda, Carmen I. Burghelea, Antonio Palanca-Soler

## Abstract

An ecosystem is generally sustained by a set of integrated physical elements forming a functional landscape unit – ecotope, which supplies nutrients, microclimate, and exchanges matter and energy with the wider environment. To better predict environmental change effects on ecosystems, particularly in critically sensitive regions such as high altitudes, it is imperative to recognise how their natural landscape heterogeneity works at different scales to shape habitats and sustain biotic communities prior to major changes.

We conducted a comprehensive survey of catchment physical, geological and ecological properties of 380 high altitude lakes and ponds in the axial Pyrenees at a variety of scales, in order to formulate and test an integrated model encompassing major flows and interactions that drive lake ecosystems.

Three composite drivers encompassed most of the variability in lake catchment characteristics. In order of total percentage of variance explained they were: (i) hydrology/hydrodynamics-responsible for type and discharge of inlets/outlets, and for water body size; (ii) bedrock geomorphology, summarizing geology, slope and fractal order-all dictating vegetation cover of catchment slope and lake shore, and the presence of aquatic vegetation; and, (iii) topography, i.e. catchment formation type-driving lakes connectivity, and the presence of summer snow deposits. While driver (i) appeared to be local, (ii) and (iii) showed gradient changes along altitude and latitude. These three drivers differentiated several lake ecotopes based on their landscape similarities. The three-driver model was successfully tested on a riparian vegetation composition dataset, further illustrating the validity and fundamental nature of the concept.

The findings inform on the relative contribution of scale-dependent catchment physical elements to lake ecotope and ecosystem formation in high altitude lakes, which should be considered in any assessment of potentially major deleterious effects due to environmental/climate change.

Lake Bassia at 2275m a.s.l. in the Pyrenees National Park, France, is one of the millions of remote high altitude lakes worldwide whose catchments are likely to experience severe effects due to climate change. Photo credit: Antonio Palanca-Soler.

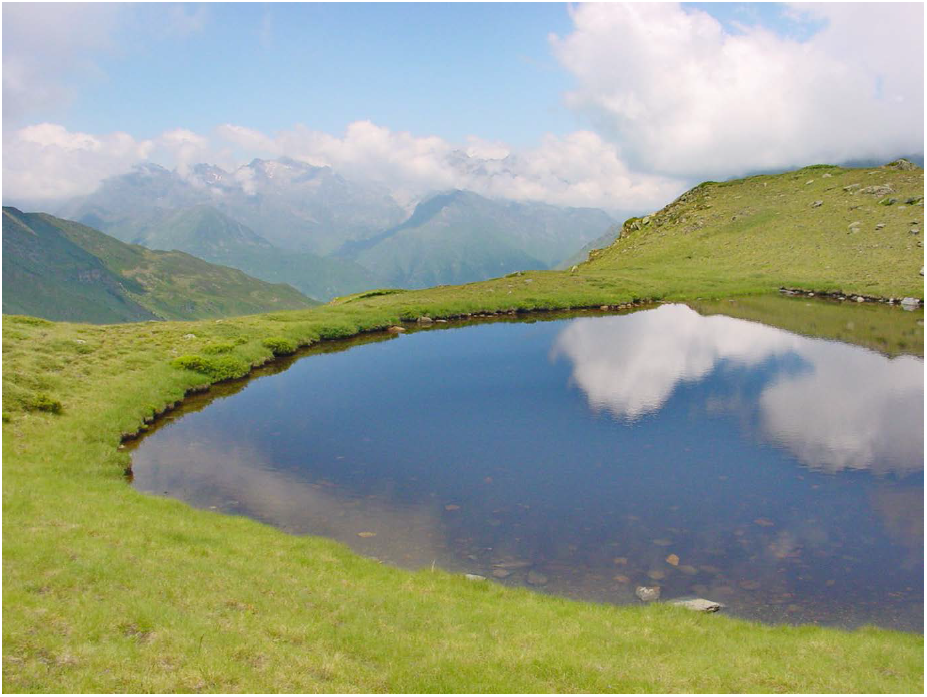

## 1. INTRODUCTION

One of the first conceptual ideas illustrating ecosystem-landscape interdependence was Vernadsky’s theory of Earth’s surface evolution, which recognized the synergetic relationships and transfer of nutrients between geosphere and biosphere (Vernadsky 1926). Recent research in the critical zone framework (i.e. Earth’s near-surface environment influenced by life; Richter and Billings 2015) advances this understanding by providing high spatial and temporal resolution details of landscape physiology at a variety of scales.

From a landscape perspective, a lake is a structural and physiological unit that draws energy and nutrients from its surrounding catchment. A lake ecosystem is therefore sustained by its physical template (ecotope, the lake’s life support system), which incorporates elements of catchment geomorphology, land cover and climate, all directly and indirectly affecting the flows of water and nutrients resulting from bedrock weathering. Predicting how changes in physical environment control ecosystems in high altitude catchments is generally challenging, due to their remoteness, the complexity of their landscape, and the many direct and indirect linkages between landscape features and processes operating at different scales. For example abiotic factors such as water resilience and cycling, primary productivity and nutrient availability are all key aquatic factors shaping community/ ecosystem development (Van der Molen and others 2003).

Of more than 300 million lakes on the Earth’s surface, a great abundance occur at mid-to-high altitudes (Downing and others 2006). In the Pyrenees, a relatively low-density lacustric region, there are an estimated 1030 lakes of > 0.05ha above 1000m altitude (Castillo-Jurado 1992), meaning that high altitude lakes mediate a great portion of ecological and geochemical processes in mountain catchments. Due to their remoteness and high topography, most of these lakes host pristine or semi-pristine ecosystems, and are under increasing attention worldwide as clean water repositories, hotspots of biodiversity (Gopal and others 2000), sensors of long-range transported pollutants (Andrea and others 2007) and global climate change (Williamson and others 2009]). Moreover, their location in headwater basins implies that they are the first to collect and redistribute bedrock-derived nutrients to the wider biosphere. These waterbodies and surrounding catchments are therefore ideal for studying how physical environments sustain their ecosystems, before climate change can induce major deleterious effects.

Environmental influence on species richness in mountain-top lakes has been discussed in the conceptual framework of Equilibrium Theory of Island Biogeography (Vuilleumier 1970; Barbour and Brown 1974; Brown and Dinsmore 1988). The theory predicts species composition at equilibrium, in a suitable habitat, being a function of habitat isolation, size and composition (MacArthur and Wilson 1963; Losos and Ricklefs 2009). For example general trends in fauna and flora functional composition can be predicted by local physical characteristics, including geology, geomorphology, waterbody size, slope and land cover (Della Bella and others 2005; Mazerolle and others 2005; Goebel and others 2006).

At any given time, a lake/pond can be assumed to support a type of vegetation and fauna whose composition is constrained by substrate and ecotope characteristics. This could result in a particular configuration of nutrient distribution, microclimate, and a local ecosystem succession/evolution in time. It is therefore critical to understand the relative contribution of the physical elements of an ecosystem to biota development, and how they connect to regional, continental and global gradients in substrate and climate. This could address a major need in ecology, to better model how physical heterogeneity within an ecosystem predicts current and future ecological dynamics, particularly in human-induced climate and habitat stress scenarios.

We will use the term “ecotope” to represent the lake/pond and its proximal catchment area as an integrated physiological unit that supports an ecosystem. Similar to spatial patches in landscape ecology (Forman 1995), we assume this unit to represent unique combinations of hierarchically organised abiotic drivers that interact and drive the flow of energy/nutrients at multiple spatial scales, ultimately feeding and shaping the development of a lake ecosystem. The ecotope concept allows considering all such features and their spatial heterogeneity, including how they may be connected to large-scale gradients in substrate and climate, and thus has the potential to incorporate and predict their function. This concept has been used variously in the scientific literature, particularly in the framework of geographic information systems (GIS) as surface ecotope patterns for environmental conservation and in human impact scenarios (Whittaker and others 1973; Van der Molen and others 2003; Yue and Li 2010; Gwata and Mzezewa 2013; Liu and Pan 2014;Sorosjinda-Nunthawarasilp and Bhumiratana 2014). However, the concept is still confusing, and we lack sufficient empirical examples that integrate the ecosystem and its underlying abiotic drivers (ecotope) in clear conceptual models/units that would better fit into the current (interdisciplinary) paradigm of Earth function as a life support system.

The main aim of this work was to identify the main landscape elements assumed to sustain a lake ecosystem in high altitude basins, and model how they organise at different scales to produce a coherent ecosystem functioning. We also postulated that a lake’s physical template is not formed randomly. Rather, it is a geomorphic inheritance left by the past major transformations of the landscape, particularly following the last glaciation. The work is based on a survey of 380 waterbodies in the axial Pyrenees. The strong E - W orientation of this mountain range, together with large blocks of distinct geology provide sharp contrasts in climate and biogeography, that makes the concept easier to test against large geographical gradients.

A secondary aim was to identify and define a number of ecotope types, supporting distinct lake ecosystems, which integrate related physical drivers. The lake ecotope concept is ultimately validated by showing its effect on lake riparian ecosystem composition.

## 2. METHODS

### 2.1 Study area and geology

The Pyrenees extend over roughly 430 km from the Atlantic to the Mediterranean Sea and separate the Iberian Peninsula from the rest of continental Europe. The area under study extends over about 80 km in the axial part of Pyrénées National Park (Atlantic Pyrenees), France (Fig. 1). This area, under reinforced protection, is restricted to recreational hiking, angling, and seasonal livestock grazing. Due to their location, the majority of the waterbodies could be considered to reflect mostly natural processes.

Bedrock geology is marked by the outcrop of Cauterets-Panticosa igneous (granitic) batholith in the central part, flanked by metasedimentary (shale) and sedimentary (limestone) materials (Fig. 1). The abundance of granite, which is particularly resistant to erosion, gives the region a characteristic steep-sloping aspect. The contact zone between the granitic outcrop and the low-grade metamorphic material includes ore deposits, some of which have been exploited for metalliferous mining in the past (Paegelow 2008). Mineral springs are abundant in this area, particularly the hot springs at the contact of granite with the stratified rocks.

**Figure 1.**
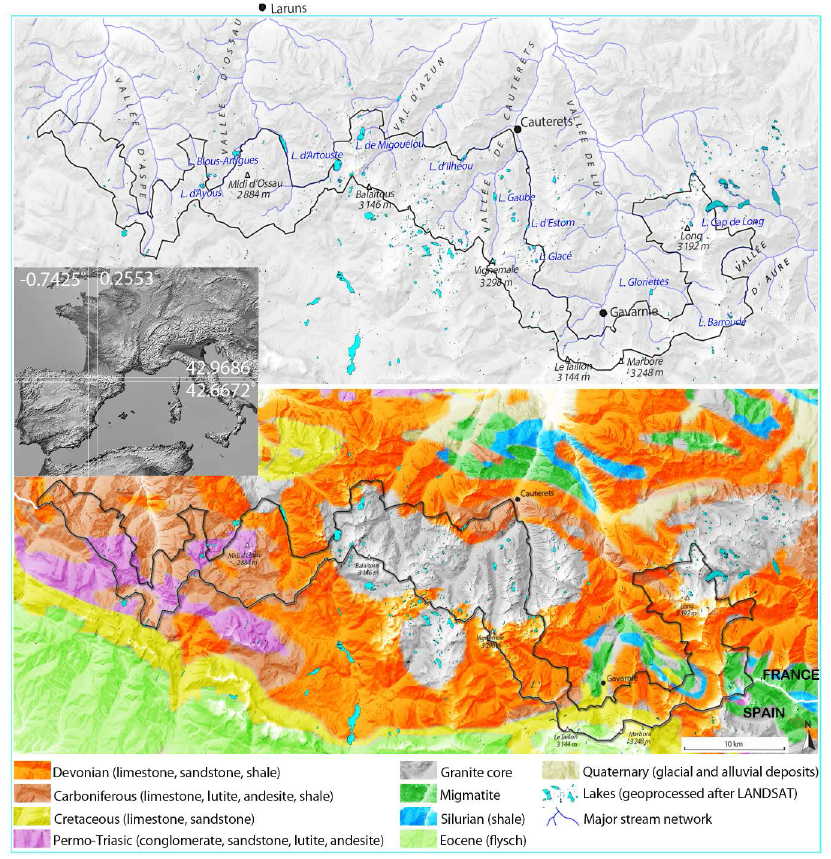
Study area (axial Pyrenees National Park, outlined in dark green) together with major hydrological and geological formations on digital elevation model. Lake representations are after LANDSAT imagery; inset radar map is after JPL (2000); Pyrenees digital elevation model is after Geoportail (http://www.geoportail.fr/); geological representation is after SGN (1996).

### 2.2 Climatology

The main air masses are from the W – NW, bringing precipitation (i.e. rain, snow and moist air) mainly from the Atlantic and the Bay of Biscay (oceanic-suboceanic climate; Mate 2002). This leads to a marked contrast between different sections/valleys of the region with glacial formations being formed mainly on the N-oriented slopes of the western and the central parts of the range. Some of the glaciers are still active and are the source of major streams. Precipitation averages 100–160cm year^−1^ in the area while mean annual temperature is 13–14^°^C (0^°^C isotherm oscillating between 1200m in January-3300m in July/August). Tree line varies between 2000-2500m a.s.l. The snow cover above 2000m settles down in November and starts to thaw in April. The glacier forming line is relatively high, ranging between 2500 to 3200m a.s.l. (Kessler and Chambraud 1990).

**Table 1.** Description of geographical and ecological variables used in the analysis of 380 altitude lakes from the central Pyrenees.

| Parameter | Values |
| --- | --- |
| Latitude<sup>+</sup> | Geographic coordinates |
| Longitude<sup>+</sup> | Geographic coordinates |
| Altitude<sup>+</sup> | Meters a.s.l. |
| Catchment type<sup>+++</sup> | Plain, U shape valley, slope, mountain pass, V shape valley, head of glacial valley |
| Main geology<sup>+++</sup> | Conglomerate-sandstone-claystone, limestone (+sandstone-marlstone-schist enclaves), schist (+andesite-sandstone-claystone and granite-limestone), granite (+schist) |
| Size<sup>++</sup> | Pool (<315±333m<sup>2</sup>), pond (1566±1985m<sup>2</sup>), small lake (9157±10267m<sup>2</sup>), medium size lake (41127±31820m<sup>2</sup>), large lake (91441±37307m<sup>2</sup>) |
| Fractal order<sup>++</sup> | 1-4 scale |
| Visible connectivity with other<sup>+++</sup> | Absent, surrounded by another lake, with another one, in chain |
| Nature of water input<sup>+++</sup> | Meteoric, spring, stream/waterfall |
| Tributary discharge<sup>++</sup> | Absent, low discharge, medium discharge, high discharge |
| Nature of water output<sup>+++</sup> | Absent, temporary, surface-small, surface-medium, surface-large, subterranean, dam output |
| Aquatic vegetation<sup>++</sup> | Absent, Absent but water flooding the grassland, scarce, abundant |
| % grass covered shore<sup>++</sup> | <10, 11-20, 21-30, 31-40, 41-50, 51-60, 61-70, 71-80, 81-90, 91-100 |
| % grass covered slopes<sup>++</sup> | <10, 11-20, 21-30, 31-40, 41-50, 51-60, 61-70, 71-80, 81-90, 91-100 |
| Slope of lake perimeter<sup>++</sup> | Plain, plain in alternation with medium slopes, medium slopes, steep in alternation with medium/plain, steep in >50% of perimeter |
| Shore snow coverage<sup>++</sup> | Absent, <10%, 11-50%, >51%, into the water |
| Snow deposits in the catchment<sup>++</sup> | Absent, very scarce, scarce, abundant, very abundant |
| *Shape of lake/pond<sup>+++</sup> | Circular, elliptic, elongate, irregular, triangular, rectangular, in 8, boomerang |
| *Modifications in lake's shape<sup>++</sup> | Absent, one input/output stream, various input/output streams |
| *Color<sup>+++</sup> | Blue-grey, opaline blue, opaline white, turquoise green |
| *Water level marks<sup>++</sup> | Absent, < 50cm, > 50cm |
| *Damming<sup>++</sup> | Absent, small dam, big dam |
| *Shore vegetation coverage<sup>++</sup> | Most of it, partially, scarce |
| *Shore coverage<sup>+++</sup> | Scarce vegetal cover (>50% cliffs, >50% slope drift, cliffs+slope drift, bedrock, bedrock+slope drift, bedrock+dispersed rocks, big granite blocks), medium vegetal cover (bedrock+grass patches, grassland+rocks, grassland+slope drift+rock blocks, cliffs+slope drift+ grassland, slope drift+grassland+scrubs, forest+cliffs+slope drift+grassland area) and dominant vegetal cover (>50% grassland, >50% scrubland, grassland+scrubs+forest, grassland+dispersed rocks, grassland+scrubs+rocks, grassland+bedrock+rocks, sheep field) |
| *Coverage of near catchment<sup>+++</sup> | Scarce vegetal cover (>50% cliffs, >50% slope drift, cliffs and slope drift), medium vegetal cover (cliffs with slope drift and vegetated patches, grassland with scrubs and rocks, grassland with cliffs and slope drift, cliffs with slope drift, grass patches, scrubs and forest) and dominant vegetal cover (>50% grass land, grassland and scrubs, forest with grass land and scrubs) |

Variable measure: <sup>+</sup> scale (numerical), <sup>++</sup> ordinal (categorical) and <sup>+++</sup> nominal. Variables proceeded by superscript (\*) did not improve PCA total variance, and were removed from analyses.

### 2.3 Hydrology

There are more than 400 lakes and ponds within the boundaries of the Pyrénées National Park. The great majority of the lakes are of post-glacial origin and they are formed at valley head, in the axial part of the mountain range. They are relatively small water-bodies, as most (>90%) of the lakes on the Earth’s surface (Downing and others 2006). A large number of streams (>210), locally called *gaves*, drain the lake catchments, and give the hydrological network a dendritic structure (Fig. 1). These ‘gaves’ subdivide the area in six major units: Aspe, Ossau, Azun, Cauterets, Luz and Aure (Fig. 1 and Appendix S1). A total of 13 lakes in our dataset were transformed into reservoirs (Appendix S1), and are used for providing fresh-water and hydropower (Mate 2002).

### 2.4. Sampling and statistical methodology

In total 380 lakes/ponds were surveyed during the month of July in 2000, 2001 and 2002. The sampling was aimed to represent the majority of mountain lakes in the area and it was undertaken in an east-westward direction to minimize potential bias due to a generally late snow thaw in the western side. Appendix S1 lists the names and locations of the surveyed water-bodies. At each site a number of major landscape factors assumed to influence ecotope processes were visually approximated and scored according to dominant units. A detailed description of the variables surveyed is presented in Table 1. Lakes’ size categories were estimated from their surface area. This was approximated as the surface area of an ellipse whose major and minor diameters were measured in the field. A digital laser telemeter was used for this purpose. Furthermore, a portable GPS device was used to record the latitude, longitude and altitude at each location.

A riparian vegetation survey was completed around each lake by visual inspection, at the same time when other environmental parameters were measured (Zaharescu 2011). The species were identified in the field using Grey-Wilson and Blamey (1979), Fitter and others (1984) and GarcÍa-Rollán (1985) keys, and where not possible they were transported in a vasculum and identified in the laboratory.

Major statistical steps are summarised in Fig. 2. Principal Component Analysis (PCA) was used as an exploratory first step to disentangle the complex relationships between the large number of landscape variables, and reduce them to a small number of variable sets, termed principal components (PCs), that can reflect major environmental drivers of a lake ecotope. Each PC is composed of related variables and it is uncorrelated to the other PCs. Since PCA is not p-value driven, the technique is robust for non-normal data distributions. It also seemed to fair better than categorical PCA on our dataset, by producing more meaningful PCs. A Varimax rotation with Kaiser normalization was applied to the extracted axes (components) in order to maximize the captured variance, while keeping the uncorrelated nature of the PCs. Variables that did not improve the model i.e. they decreased the factor total explained variance, were excluded (Table 1). For this analysis name variables (e.g. geology) were re-coded on ordinal scales representing major transitional phases (e.g. from sedimentary, to metamorphic, to igneous bedrock) before using them in PCA. Results of PCA were interpreted using this encoding. The Kaiser-Meyer-Olkin measure of sampling adequacy in PCA was 0.72, and the Bartlett’s test of sphericity, approx. *χ*^2^=1398.2 (P<0.001), indicating that the analysis was reliable and adequate.

**Figure 2.**
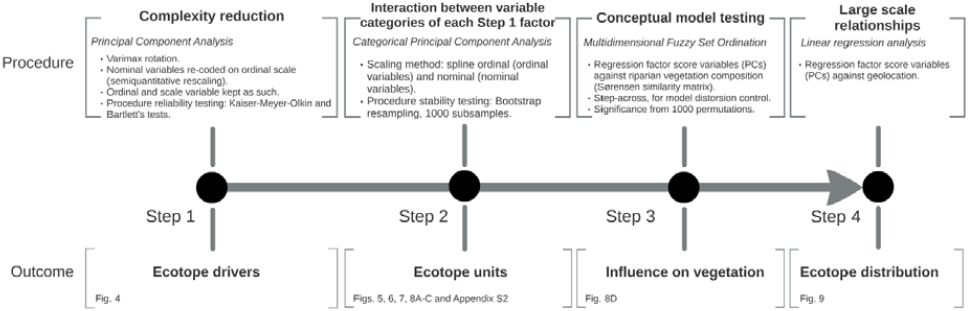
Flow chart illustrating the sequence of statistical steps supporting this work, and their outcomes.

To help identify and exemplify ecotope units, the spatial interaction between categories of variables in each extracted factor in step 1, and their vector projection on lakes in the ordination space were evaluated using categorical principal component analysis (CATPCA; Fig. 2). This is a nonparametric approach capable of finding relationships between a wide range of nonlinear variables (including numerical, categorical and nominal variables) and between variable categories and cases (lakes). Because CATPCA stability may be influenced by the sample characteristics, including sample number and scaling method, the degree of sensitivity to changes in the data was tested by bootstrap resampling. This procedure used 1000 random sets of subsamples from the original dataset and repeated CATPCA on each set. It determined the constancy of assignment (correlation) of the variables to the component vectors by producing 90% confidence regions of component loadings. If the results provided by CATPCA are stable, we expect narrow confidence ellipses.

The reliability of ecotope factors was tested for their influence on riparian vegetation composition using the logic of Multidimensional Fuzzy Set Ordination – MFSO (Roberts 2008). This approach related the explanatory PCA-derived composite factors (summarized into regression factor score variables) to riparian vegetation structure (response variables) by using a distance matrix of species incidence (calculated using S0rensen similarity index). Generally, this matrix gives a measure of similarity between sites based solely on biota composition. Multidimensional fuzzy set ordination is a more natural alternative to classical ordination methods based linear algebra logic, which assume objects to either belong (1) or not (0) to a given set or function. Instead, MFSO assumes that a case (element) can have partial membership values between 0 and 1, thus creating a range (fuzzy) of influences into the model. Since ecosystem-environment interactions are not always restricted to well-defined algebraic functions (they can be discontinuous), MFSO is particularly suitable to solving partial relationship problems, which are more characteristic to real-world phenomena. Likewise, in MFSO factors have to be chosen beforehand and their contribution to the model is independent, therefore hypothesis testing of ecological/environmental processes (e.g. cause-effect relationships) is possible, for instance in gradient analyses (Roberts 1986). To improve the distortion effect given by sites with no species in common, a step-across function was used together with MFSO (Boyce 2009). This approximation procedure finds dissimilarities between sites (cases) above a threshold of missing data (no species in common) and replaces them with the shortest paths by stepping across intermediate sites. Visually, it produces an expansion of the data cloud at the distortion site (matrix dissimilarity before the procedure=1), improving therefore the fit. The significance of MFSO model was drawn after 1000 permutations.

**Figure 3.**
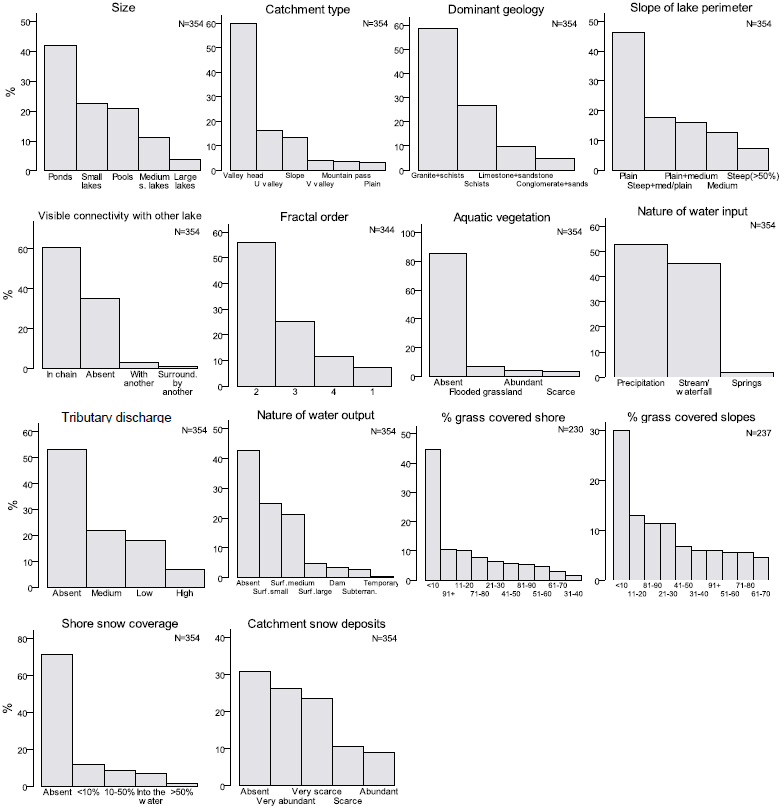
Frequency distribution (%) of sampled landscape variables in 380 altitude lake/pond locations from the Central Pyrenees.

As a final step, linear regression was used to examine the potential relationship between catchment-scale ecotope properties and large scale geographical gradients (Fig. 2). The variables were summarised as regression factor scores of the extracted principal components (PCs) before being used as response variables to geographical predictors in the regression analysis. Statistical treatment of the data was conducted in PASW for Windows (former SPSS, SPSS Inc. 2009). Bootstrap procedure was computed with macro file Categories CATPCA Bootstrap for PASW developed by Linting et al. (2007), available online at http://www.spss.com/devcentral/. Multidimensional Fuzzy Set Ordination was computed in R statistical language, using FSO (Roberts 2007) and LabDSV (Roberts 2012) packages. Step-across function was performed in VEGAN package (Oksanen and others 2012) for R (R Development Core Team 2005).

## 3. RESULTS AND DISCUSSION

The surveyed water-bodies spanned from 1161 to 2747 m altitude. Figure 3 presents the frequency distribution of the assessed landscape variables. As can be observed from this figure, most of the water-bodies can be included into pond and small lake categories. These waterbodies are mostly located on relatively flat surfaces at the head of glacial valleys; they have granite-dominated bedrock, and a great number of them are connected in chain with other lakes/ponds within their area. Likewise, the central Pyrenees lakes showed weakly developed riparian zones (as indicated by a high frequency of lakes with low fractal order, Fig. 3), which correspons to a relatively young age on a lake evolutionary time scale. With few exceptions aquatic vegetation was largely absent at the time of sampling. Most of the lakes/ponds are fed by precipitation or small surface streams of very low discharge, which is typical of high altitudes. Accordingly, a great number of them have visibly absent or small outputs. Water flowing from springs, on the other hand, seemed to have very little importance in their hydrodynamics. Shore/slopes vegetation coverage for most of the water-bodies was < 10%, and a mixed snow coverage was recorded in their near-catchment during the survey.

### 3.1 Deconstructing the main drivers of a lake physical template

The interaction between climate and geomorphology can potentially shape the formation of ecotopes. To examine the influence of catchment-scale landscape components on the structure of lake ecotopes, a PCA of all assessed variables (Table 1) was carried out. This reduced the variables to a limited number of key components (composite factors) which can explain the main environmental drivers of lake characteristics. The inherently complex nature of the high altitude environment meant the total PCA variance in our dataset was split over >3 uncorrelated composite factors (Fig. 4 inset). The first three components (PC1, PC2 and PC3) together accounted for more than 58% of the total variance in the lake and catchment characteristics (Fig. 4). The first component (PC1) accounted for 21.3% of the variation (Fig. 4). It, i.e. PC1 (interpreted hereafter as hydrodynamics), indicates a strong association between waterbody size and lake hydrology (type and volume of water input/output). This is important as aquatic macrophytes and invertebrates richness are likely to vary with the size of a lake (Oertli and others 2002; Biggs and others 2005), a core idea in the “ecological theory of island biogeography” (MacArthur and Wilson 1963; Losos and Ricklefs 2009).

**Figure 4.**
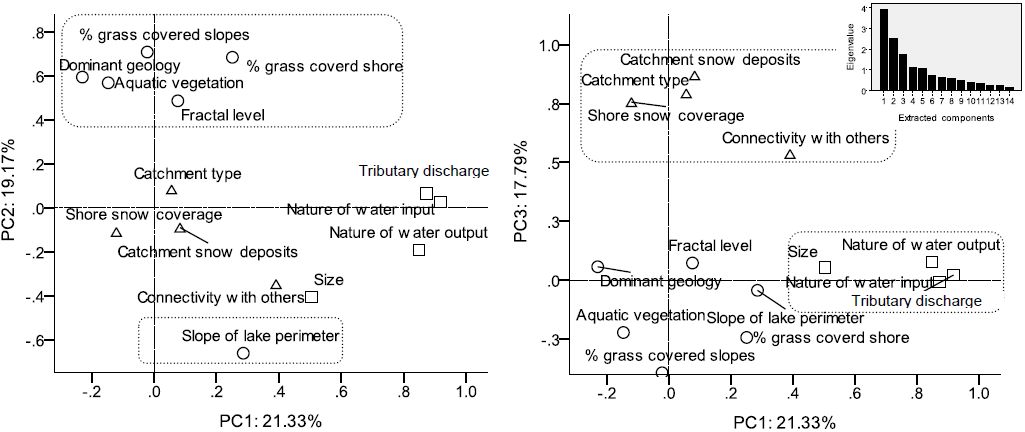
Relationships between landscape variables in their projections on principal components 1–2 and 1–3, of principal component analysis (PCA). Variables clustering with the PCs are enclosed in dash line. Figure symbols represent the variables with high loading on: (□) PC1, (○) PC2 and (Δ) PC3. Inset plot shows the number of extracted components. N=380 water-bodies.

The second component (PC2, explaining additional 19.2% of the total variance) had high loadings for the variables that would be determined by the main bedrock geology/ geomorphology, i.e. geology, shore sloping, % of slope/shore covered by grass, fractal order and the presence of aquatic vegetation (Fig. 4). Geerling et al. (2006) have shown that ecotope composition (riparian surface, vegetation coverage and composition) can change during rejuvenating hydro-geomorphological processes of rivers, i.e., meander progression, meander interruption and channel shift. Likewise, substrate geology and slope are recognised physical factors that can influence the characteristics of a lake through their effects on hydraulics, weathering and nutrient cycling processes which together shape its biological structure (EC 2000; Kamenik and others 2001). It seems therefore that geo-morphology is a second major driver of an altitude lake ecotope development and can influence not only the topographically-related high energy processes, such as slope erosion and runoff, but also the riparian development, its vegetation coverage and the development of aquatic vegetation. Lake shores’ vegetation coverage is a crucial ecotope factor in high altitude waterbodies which has been found to control nutrient cycling in a lake and therefore its biotic composition (Kopacek and others 2000).

Finally, the third PC axis accounted for further 17.8% of the variability in the lakes’ characteristics. The variables grouped under PC3 were: presence of snow deposits at shore level and in the near catchment, catchment type and visible connectivity with other lakes, together being interpreted as topographical formation (Fig. 4). The PC3 findings suggest that topography also has significant control over ecotope processes by its influence on important factors such as habitat connectivity and habitat snow coverage, the latter being important in shaping land-water interactions during the large periods a mountain lake catchment is snow-covered (Edwards and others 2007).

**Figure 5.**
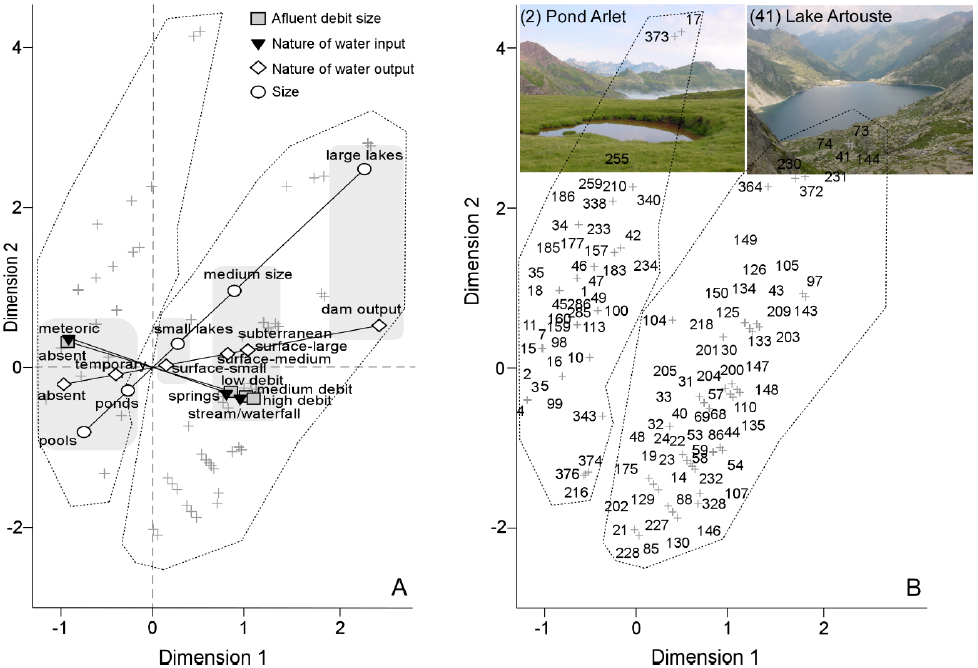
(A) Interaction between variable categories in hydrodynamics factor (i.e. PC1 in Fig. 4) and their association with lakes, as shown by CATPCA. To aid in interpretation, the associated categories were enclosed in grey, while their association to resulted lake groups is enclosed in dashed polygons. Lake grouping is further detailed and illustrated in (B). Lake coding corresponds to Appendix S1.

Evidence has shown that the patterns of snow distribution in rugged alpine terrain are the most visible consequence of topography and its interaction with climatic variables like precipitation, solar radiation and wind (Körner 1992, 2003; Gottfried and others 1999). The seasonal cycles of snow accumulation and ablation as well as snow coverage can have a crucial influence on high altitude ecosystems’ composition at a variety of scales, with species capable of coping with the environmental conditions/stresses becoming more abundant (Walker and others 1993; Keller and others 2005). Habitat connectivity, on the other hand, is an important factor in maintaining the integrity of metapopulations of plant (Biggs and others 2005) and animals (Richards-Zawacki 2009), with species assemblages likely to be richer in areas that facilitate propagule dispersal and colonisation. This is a second important aspect of Island Biogeography Theory (MacArthur and Wilson 1963) which predicts an increase in species number with a decrease in remoteness of an island ecotope.

The remaining 42% variability in the dataset is accounted for by other numerous factors, individually each accounting for a small amount of the variability (Fig. 4 inset).

**Figure 6.**
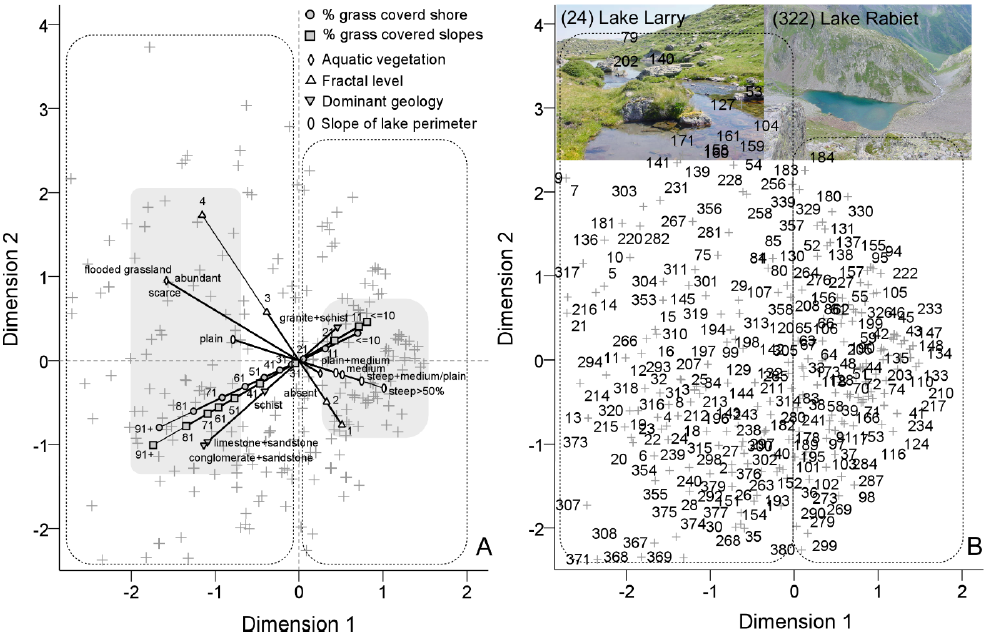
(A) CATPCA interactions between variable categories of the second extracted component (geomorphology in Fig. 4), and their association to lakes. Category groups are highlighted on grey and lake groups are surrounded by dashed polygons. (B) Grouped lakes, with coding detailed in Appendix S1.

### 3.2 Integrated physical drivers determine lake basin types

Further analysis of the three PCs, individually, can help uncover the influence they have on ecotope development. To classify the waterbodies into ecotope types we studied the interaction between the variable categories within individual PCs previously discussed, i.e. hydrodynamics (PC1, Fig. 5), geo/morphology (PC2, Fig. 6A and B) and topographical formation (PC3, Fig. 7A and B). The analysis yielded a considerable degree of stability, as shown by relatively narrow 90% confidence ellipses of the bootstrap component loadings (Appendix S2), and can therefore be used confidently.

As displayed in Figure 4A, the interaction between hydrodynamics variables (PC1) shows that small waterbodies such as pools and ponds are fed principally by meteoric water, e.g. snow and rain, and such water-bodies either lack or have temporary tributaries/outputs. They represent a lake ecotope category. A second category is represented by small and medium-size lakes. They are characterized by various forms of water input, including springs and streams/waterfalls of low to high discharge; this category is also associated to a diverse output nature, e.g. surface and subterranean (Fig. 5A). On the other side, large lakes plot further apart and are represented by dam lakes (Fig. 5A). The analysis also shows the cross-point where major lake properties change, with variable vectors plotting onto two well-defined waterbody clusters: the first cluster, pools and ponds of low water turnover, plotting on the negative side of the first dimension. And the second cluster, represented by small to large lakes of a relatively large tributary/output, plotting on the positive side in the ordination space (Fig. 5B). This is an important finding since waterbodies which receive significant runoff can have different biotic composition compared with the mainly rain-fed ones, as they can receive more nutrients from the catchment (EC 2000; Kamenik and others 2001). Riera et al. (2000), Saros et al. (2005) and Robinson and Kawecka (2005) provide illustrative examples of how nutrient availability/drainage type can shape phytoplankton, crayfish and fish development in oligotrophic alpine lakes.

**Figure 7.**
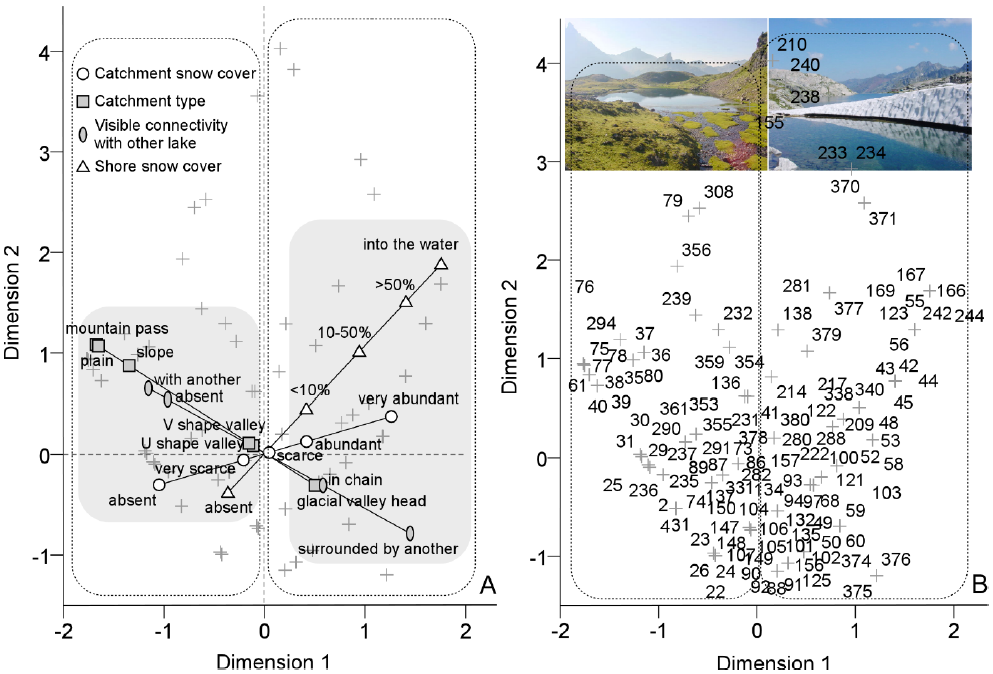
(A) Interactions (CATPCA) between variable categories of the third principal component (topographical formation, Fig. 4) and their associations to lakes. In grey are associated categories, while dashed polygons highlight resulted lake groups. (B) Lake grouping, with lakes listed in Appendix S1.

The plot of interaction between PC2 variables, representing geo/morphological processes, shows that bedrock categories such as limestone/sandstone/conglomerates associate with lakes surrounded by a relatively flat topography, >50% grass covered shore/slopes, a highly developed riparian zone and the presence of aquatic vegetation (Fig. 6A). On the other hand, granite-schist bedrock plots together with medium to steep lake shore slopes, <20% grass covered shore/slopes, a poorly developed riparian zone and lack of aquatic vegetation (Fig. 6A). These two lake categories, i.e. formed on limestone and granite, point out to a spatial segregation of lake ecotopes according to the two main geomorphological units in the Pyrenees. That is, the Paleozoic-Mezozoic sedimentary bedrock and the metasedimentary-igneous outcrops (Fig. 1) which influence biota settling at these sites. Plotting of the surveyed sites by cluster analysis, however, did not form well-defined groups, suggesting rather transient ecotope features between the two main categories (Fig. 6B), possibly owing to the influence of mixed geological compositions.

**Figure 8.**
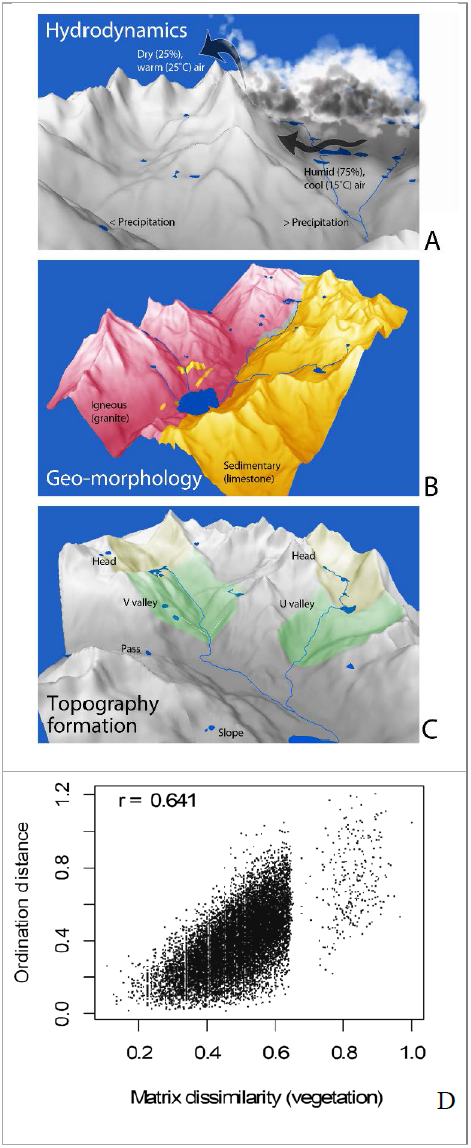
Conceptualization of lake ecotope development at high altitudes and its principal drivers, i.e. hydrodynamics (A), geo-morphology (B) and topography (C). A digital elevation model of upper Tena Valley (catchment of Lake Respomuso), central Pyrenees exemplifies this concept. The hydrodynamics gradient is represented here as a typical mountain Foehn cloud leaving the precipitation on one slope, then dissipating as it comes in contact with warm and dry air masses on the opposite slopes. Colors in geo-morphology and topography models represent distinct bedrock geology and glacial valley sections, respectively. A test of the conceptualized model performed by Multidimensional Fuzzy Set Ordination (D) shows the joint effect of the 3 ecotope drivers on the riparian vegetation species composition recorded at each lake. The number of permutations in this model = 1000.

An analysis of the third composite factor, i.e. topographical formation (PC3; Fig. 7A and B) reveals two major ecotope forms. On one hand, there are lakes at the head of glacial valleys. These are generally either interconnected in chain with other lakes, or are in a basin in the vicinity of a major lake, and have a high proportion of summer snow deposits on their shores/near-catchment (Fig. 7A). Secondly, there are lakes on flat terrain, mountain passes and V/U shaped valleys, which are generally isolated or connected to a neighbouring lake, and have very scarce or no summer snow cover in their surroundings (Fig. 7A). The geomorphic changes resulted from the last glaciation and their location in the landscape (through the extent of their influence) are likely the main drivers of this factor. This is supported by Riera and others (2000) who found differences in biota assemblages inhabiting different geomorphic settings in the Northern Highland Lake District, Wisconsin, USA.

Our analysis helped individualise and classify key physical drivers in terms of their influence on lake ecotope development. A conceptualised form of the analysis’s outcome was used to further simplify ecotope processes/forms that may be used to assess the relationship between key ecotope drivers and ecosystem functioning, e.g. vegetation structure (see below).

### 3.3 Conceptualisation and testing of a lake ecotope and its drivers

A conceptualization of the three major ecotope factors, i.e. hydrodynamics, geomorphology and topography is presented in Fig. 8. This figure illustrates different types of climate, geology and topography of the mountain terrain that are responsible for the development of different ecotopes. For example, differences in precipitation received by two slopes of a mountain as a result of Foehn cloud formation – typical of high altitudes (Fig. 8A), influence the amount of water that a lake receives as a result of a sharp drop in air moisture and an elevation of the cloud as it meets dry, warm air masses from the opposite slopes. This is a typical phenomenon found along the N-S (wet, Atlantic-dry, Mediterranean) climate gradient in the Pyrenees. Similarly, contrasting differences in substrate geo/morphology, e.g. limestone and siliceous, are fundamental for lake ecotope development (Fig. 8B). A conceptualisation of the third composite factor, i.e., topography formation (Fig. 8C) shows in a simplified way that different topographical forms or glacial formations underlie different lake types. Such influence of hydrodynamics, geomorphology and topography on ecotope formation, in *sensu amplo*, have been exemplified for other water and terrestrial environments by Van der Molen and others 2003; Hong and others 2004.

This conceptual model was tested on a dataset representing a complete survey of riparian vegetation composition in the study area (Zaharescu 2011). The effect magnitude of the three identified principal ecotope drivers on riparian plant species composition is cumulatively presented in Fig. 8D. The figure shows that with increasing vegetation similarity between sites (or decreasing dissimilarity), the environmental variability, as predicted by vegetation composition, decreases and sites become more similar in terms of their ecotope composition. There are also a fair number of sites with no species in common (the discontinuity in the data cloud), which is expected given the contrasting combination of climate, topography and geology of the study area. This means that the three drivers (PCs) identified and conceptualised in this work had a strong cummulative influence in determining the riparian vegetation species composition (cumulative Spearman *r*=0.64, *p*<0.05). The strongest influence on vegetation came from the composite factor topography formation (*r*=0.43); this was independently followed by hydrodynamics (*r*=0.12) and geo-morphology (*r*=0.09). These results imply that the conceptualised model is based on a valid identification of the key ecotope forming factors and their influence in determining ecosystem development. The approach thus has the potential to be used as a tool to predict the response of flora, fauna or other ecosystem components to changes in their life support environment.

**Table 2.**
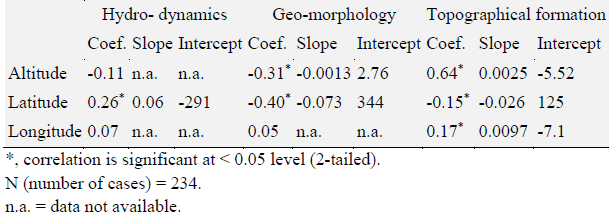
Spearman rank correlation coefficients (Coef.) between geo-position variables (as predictors) and summarised landscape variables (i.e. regression factor scores of principal components) resulting from PCA, together with regression slope and intercept values for the significant correlations. PCA factors represent: hydrodynamics, PC1, geo-morphology, PC2 and, topographical formation, PC3. Variables summarised by these composite factors are presented in Figure 3.

|  | Hydro- dynamics |  |  | Geo-morphology |  |  | Topographical formation |  |  |
| --- | --- | --- | --- | --- | --- | --- | --- | --- | --- |
|  | Coef. | Slope | Intercept | Coef. | Slope | Intercept | Coef. | Slope | Intercept |
| Altitude | -0.11 | n.a. | n.a. | -0.31* | -0.0013 | 2.76 | 0.64* | 0.0025 | -5.52 |
| Latitude | 0.26* | 0.06 | -291 | -0.40* | -0.073 | 344 | -0.15* | -0.026 | 125 |
| Longitude | 0.07 | n.a. | n.a. | 0.05 | n.a. | n.a. | 0.17* | 0.0097 | -7.1 |

\*, correlation is significant at < 0.05 level (2-tailed).
N (number of cases) = 234.
n.a. = data not available.

### 3.4 Connection with large geographical gradients

The three ecotope forming factors while fundamental to ecotope development, they may follow large scale gradients of altitude, latitude and longitude. An analysis of hydrology/hydrodynamics, PC1, geo-morphology, PC2, and topographical formation, PC3 (Fig. 4) in concert with altitudinal, latitudinal and longitudinal (macroclimate) gradients (Table 2) identified elevation as a primary gradient explaining lake ecotopes development, with local effects of the variables associated with topography, i.e. PC3 (Fig. 9A). Altitude is a geographical constraint with known influences on catchment development through its main effects on glacial processes such as cirque and valley formation. This can influence water and nutrient cycling and photosynthesis, and can lead to biota compositional differences along aquatic gradients, for example cryon/crenon – rhithron – potamon. Such examples of altitudinal effect on biota composition have been reported for various taxa, including zoobenthos, macrophyte and amphibian species (Hinden and others 2005).

Latitude was the second most important (broad-scale) gradient for lake ecotope variation, with local effects of variables related to bedrock geo-morphology (regression factor score of the second PC) (Fig. 9B). Latitude apparently also had some effect on the variables associated to lake hydrodynamics, as shown by its relatively weak, but significant relationship with PC1 regression score (Spearman ρ=0.26; Table 2). The association of latitude to geological constraints is potentially reflective of some major N-S geomorphological gradient involved in lake ecotope development. However, the variation in lake hydrodynamics across latitude could be explained by the rates at which the catchments receive moisture-charged air masses from the Atlantic Ocean along a N-S direction, which lose moisture as they advance toward the (drier) axial part of the mountain range.

**Figure 9.**
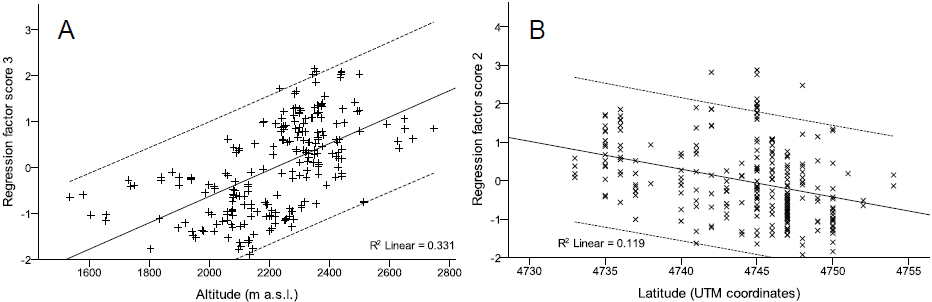
(A) Linear relationship between topographical formation (i.e. regression factor scores of PC 3: catchment type, connectivity with other lakes, catchment and shore snow coverage – see Fig. 4) and altitudinal gradient. (B) Relationship between geo-morphology (i.e. regression factor scores of PC 2: dominant bedrock geology, % grass covered slopes, % grass covered shore, aquatic vegetation and fractal development) and latitude. Confidence intervals (95%) are dashed.

## 4. CONCLUSIONS

In headwater basins of the central Pyrenees, the development of a lake ecosystem’s physical support (ecotope) was scale dependent, and was driven primarily by basin’s hydrodynamics, seconded by geo/morphology and topographical formation. These major drivers resulted in a number of lake types, which shared similarities in their catchment physical properties, and provided distinctive abiotic settings for riparian plant communities.

Except hydrodynamics, which appeared to be mostly a local factor, the identified drivers were connected to large-scale geographical gradients, of which altitude and latitude were the most influential. The relationship between a lake ecosystem and its physical template is therefore expected to change along large horizontal and vertical gradients, in connection to major substrate units, and continental-to-global climate gradients. Changes in climate factors may therefore affect not only lake ecosystem composition, as previously shown, but also many of its physical and chemical processes, such as water energy and weathering, that feed and shape fauna and flora development and their structure. Our work provides compelling empirical support of these cross-scale linkages in remote natural catchments. We interpret this as confirmation of local-to-large scale landscape evolution in the postglacial period starting 11,000 years ago, which created the major elements of the physical landscape that drove biota settling and diversification.

We conceptualised and successfully tested how hydrodynamics, geo/morphology and topography interact to support ecotope and riparian vegetation composition development. Our conceptualised template could be a common feature in mountain ranges, therefore providing an integrated and fundamental conceptual framework for hypothesis testing and experimentation in ecological modelling studies where scale and landscape properties and fluxes are important.

## ACKNOWLEDGMENTS

The study was developed with the financial support of Pyrenees National Park, France. The field work was supported over three years by an international team from Spain, UK, Romania and New Zeeland, including the late Richard Lester, as well as Javier Fernandez-Fañanas, Cristina Castan-Lanaspa, David Rodríguez-Vieites, Manuel Domínguez-Rey, Ana Quintillán-Cortiñas, Belén Cirujano-Díaz, Roberto Garcia-Carrera, Jorge Diez-Dieguez, Jorge Rodriguez-Vila, Nicolas Palanca-Castan, Claudia Toda-Castan, Jesús Giraldez-Moreira, Juan Fernández-Rodríguez, Catalin Tanase, Andreea Vasiloiu, Carles Roselló-Vila, Carlos Tur-Lahiguera, Nuria Marti, Maria José Ferrus-Leiva, Julio Palanca-Cástan, Bruce Dudley and José Martín-Gallardo – they are all gratefully acknowledged.

## AUTHOR CONTRIBUTION

Sampling campaign design, A Palanca-Soler; data collection, A Palanca-Soler and DG Zaharescu; study design and data analysis, DG Zaharescu; manuscript preparation, DG Zaharescu, PS Hooda and CI Burghelea.

## Appendix S1:Lakes Surveyed in This Study

Lakes and ponds from the axial area of Pyrénées National Park (France) analysed in this study. Altitude is in m a.s.l.; latitude and longitude are in decimal coordinates. Main valleys, locally called *gaves*, give structure to the topography.

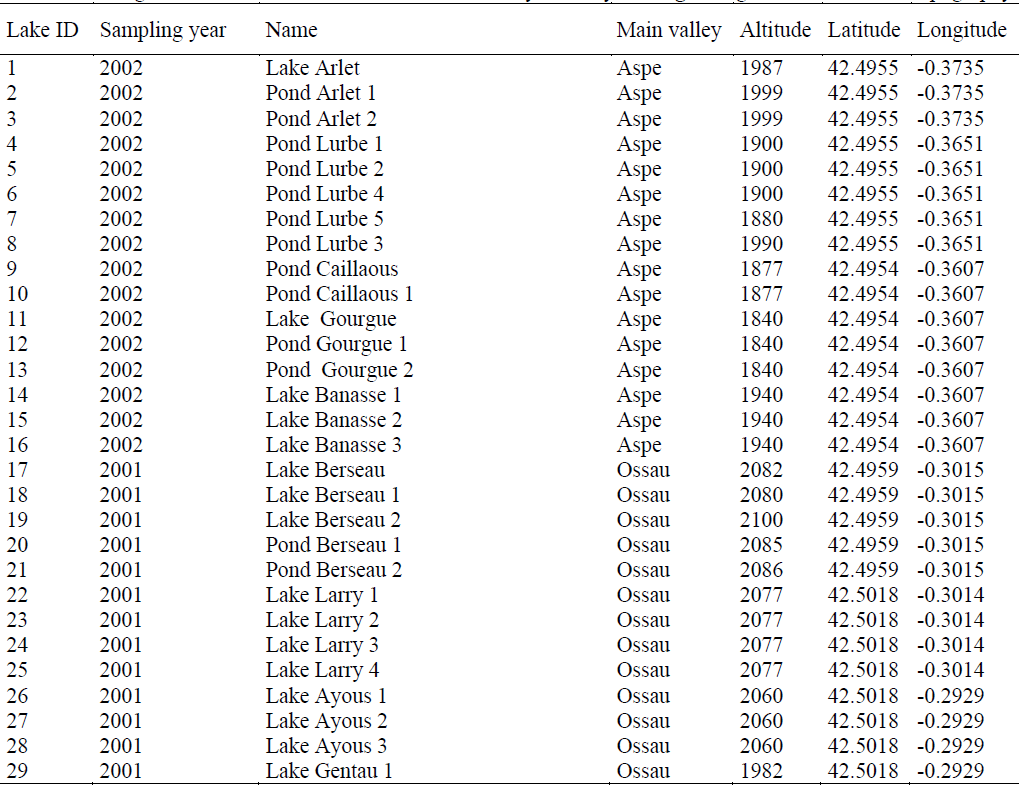

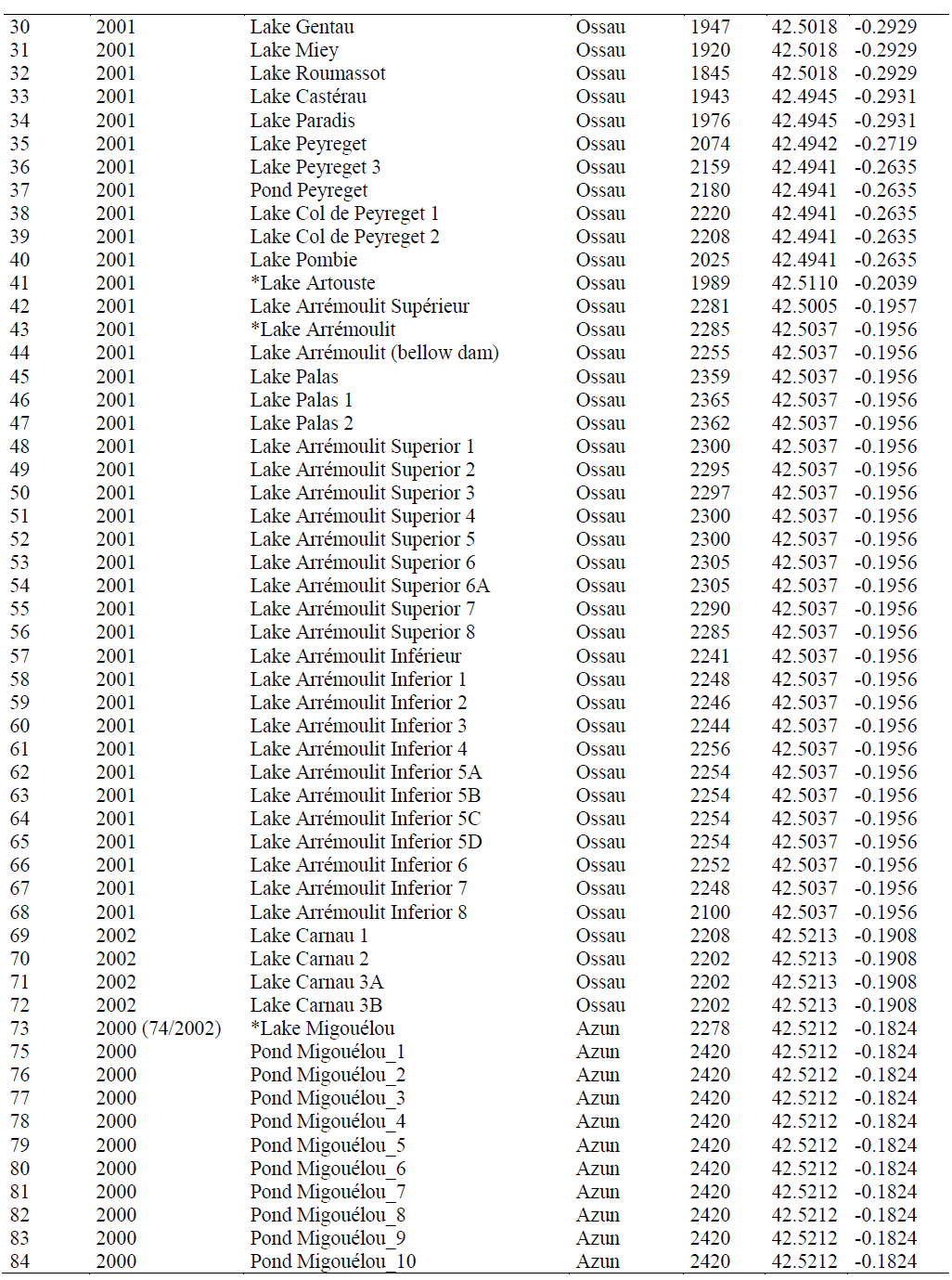

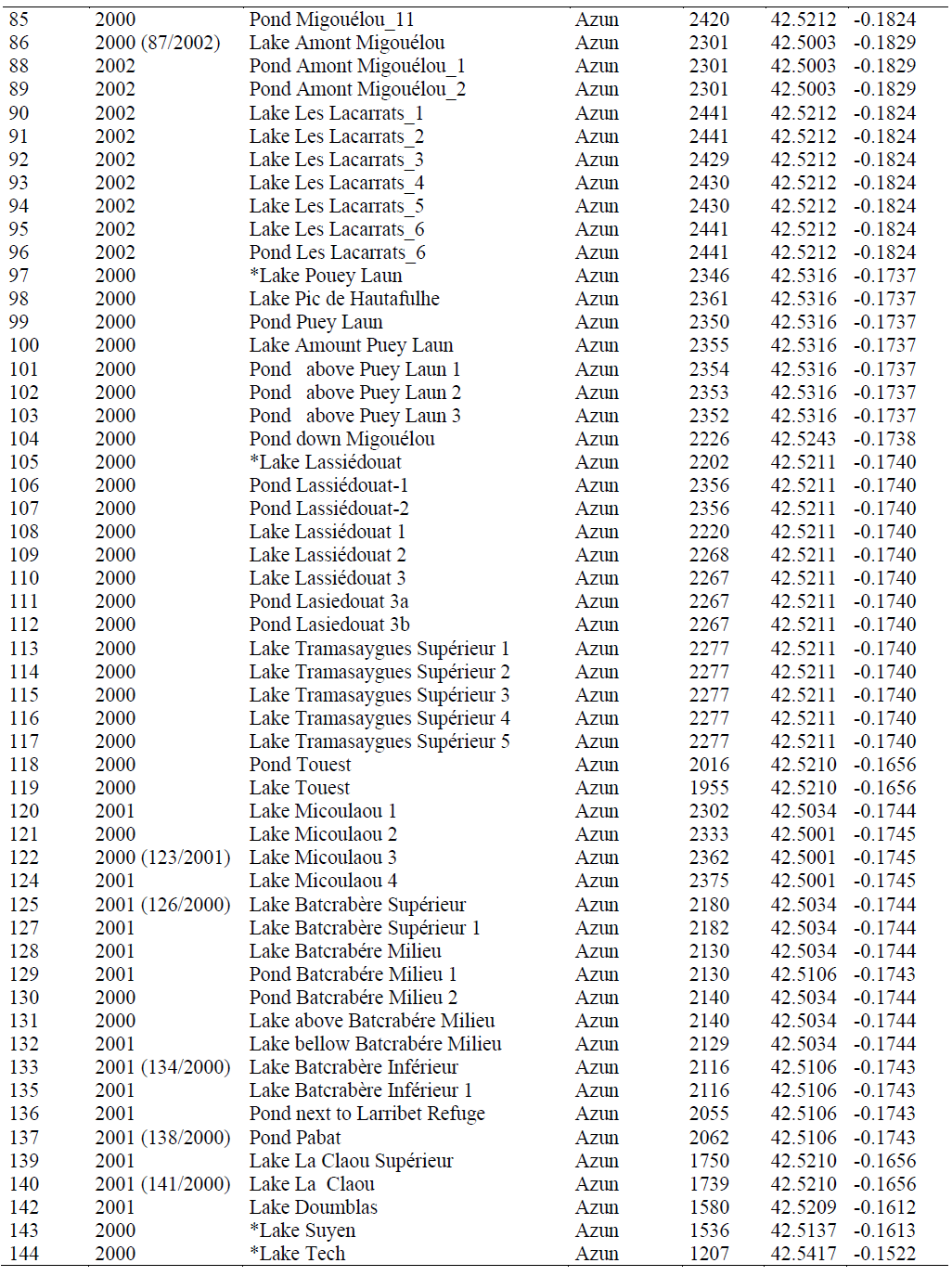

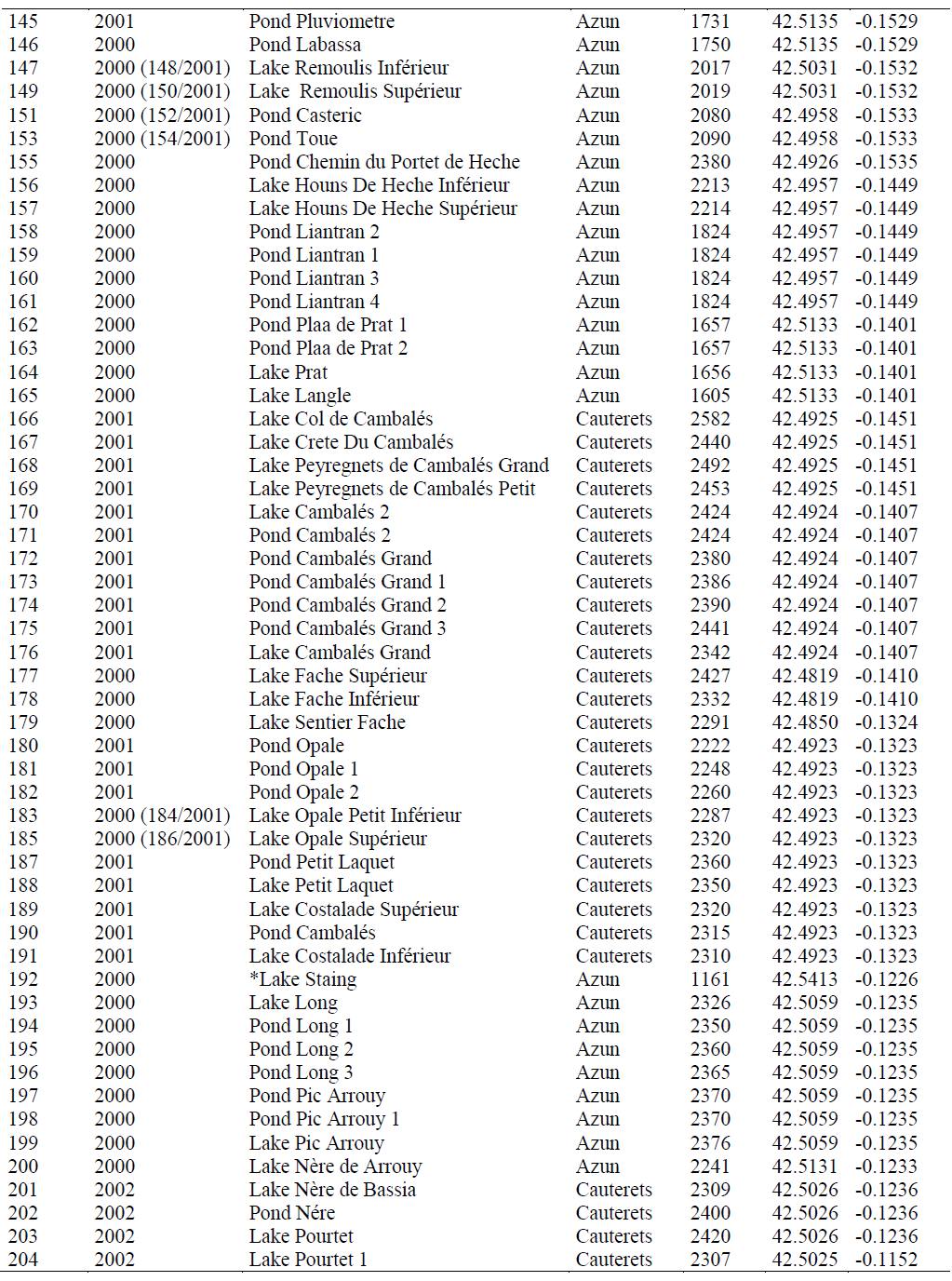

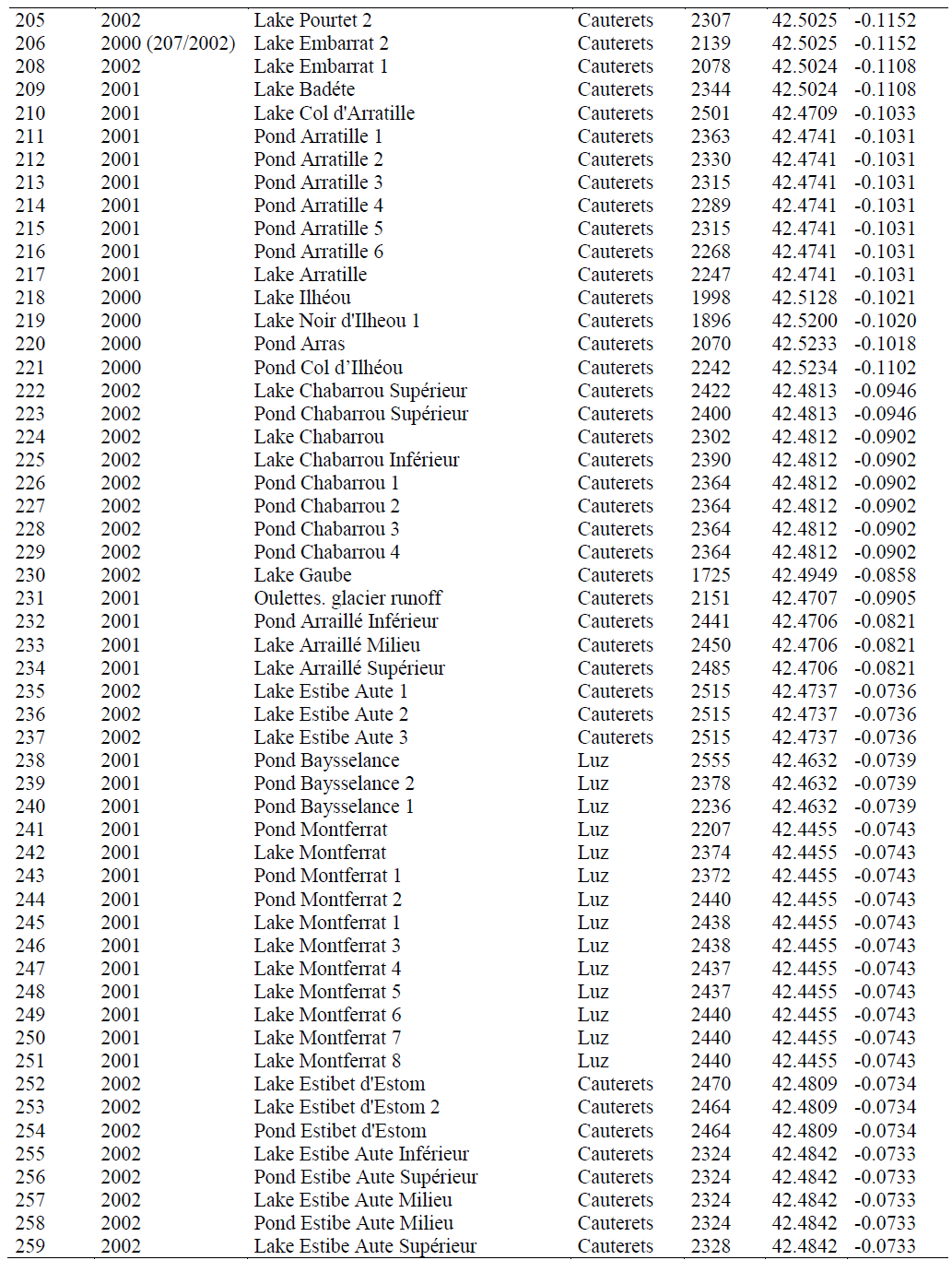

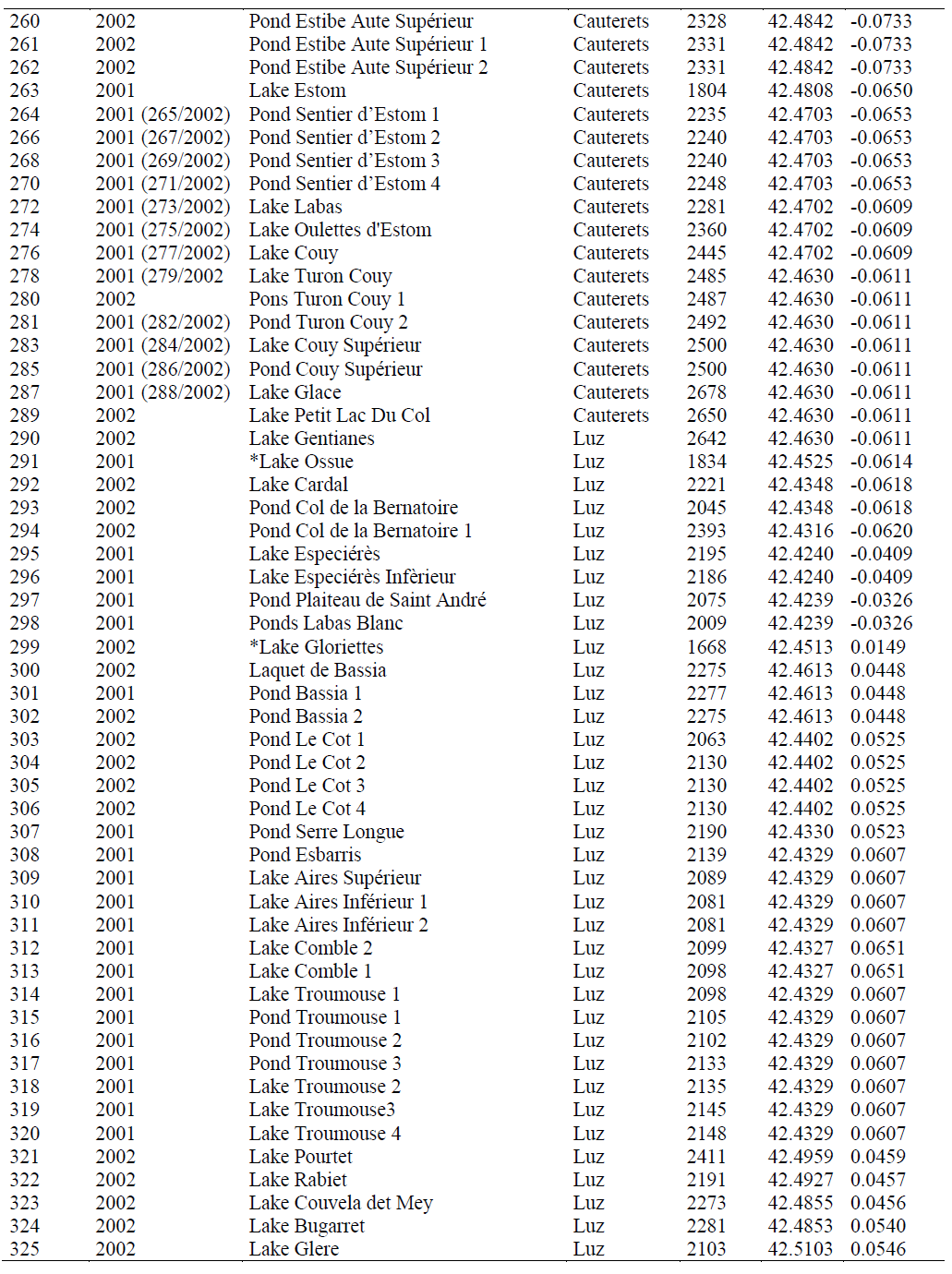

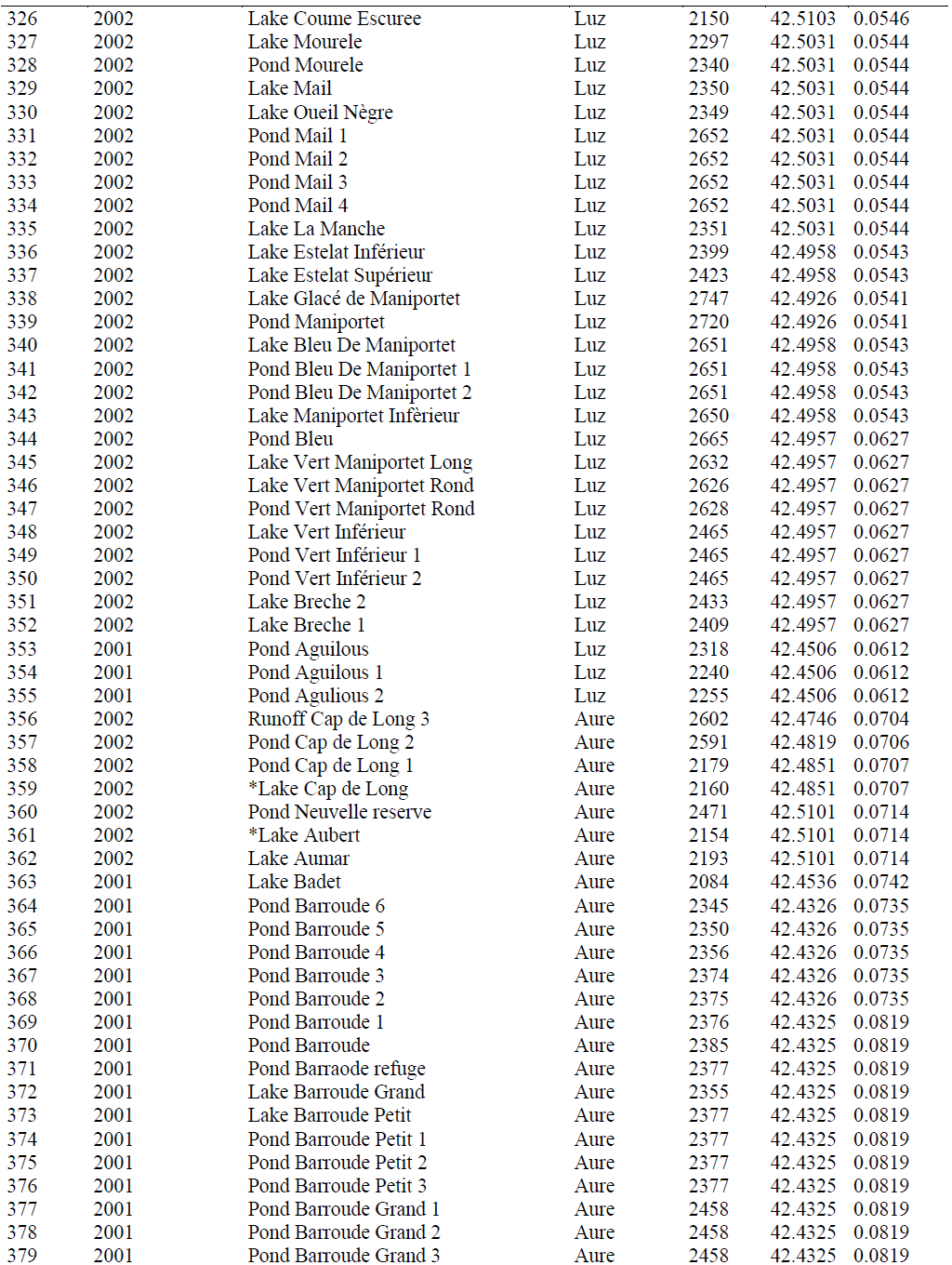

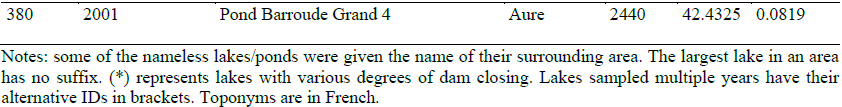

## Appendix S2:CATPCA Model Testing

Plots showing the stability of CATPCA results (i.e. variables loading on first 2 extracted dimensions), for hydrodynamics, geo-morphology and topography factors (as summarized by PCA), as given by Bootstrap resampling. Component loadings are displayed together with their 90% confidence intervals. The procedure shows a good level of stability, which is illustrated by generally narrow confidence intervals

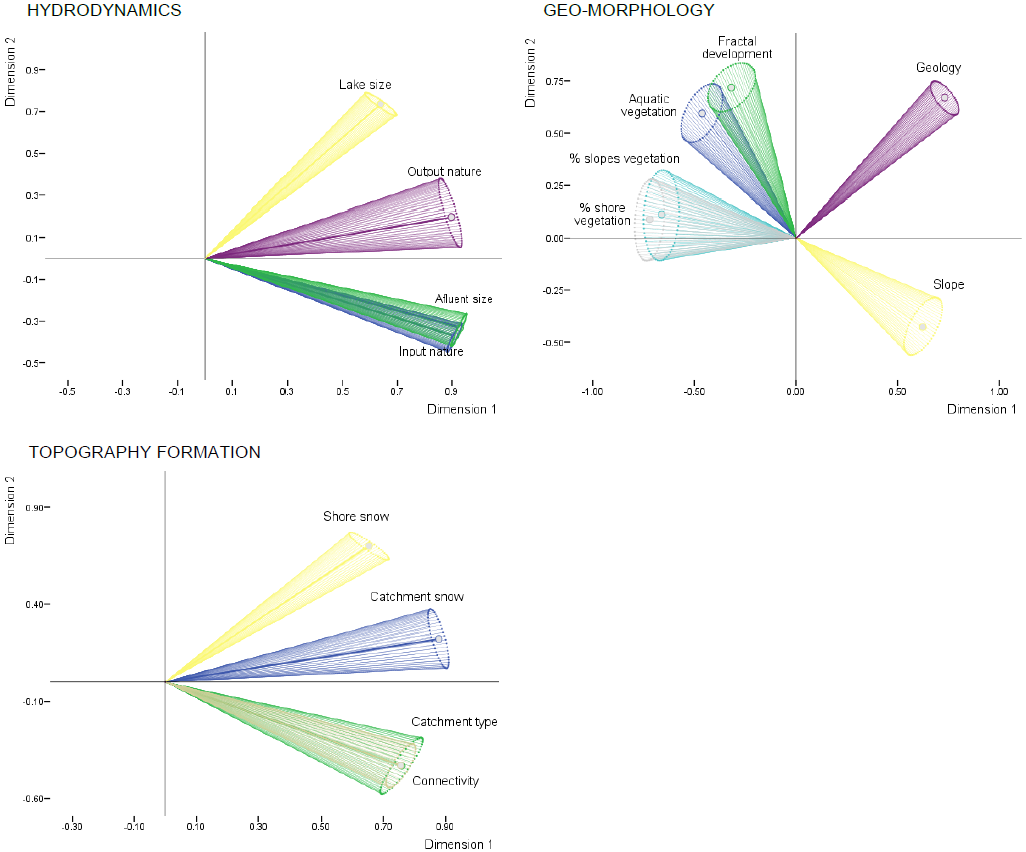

## REFERENCES

AndreaL, TartariGA, MusazziS, GuilizzoniP, MarchettoA, MancaM, BoggeroA, NocentiniAM, MorabitoG, TartariG, GuzzellaL, BertoniR, CallieriC. 2007. 21 High altitude lakes: limnology and paleolimnology. Developments in Earth Surface Processes 10:155–70.

BarbourCD, BrownJH. 1974. Fish species diversity in lakes. The American Naturalist 108:473–89.

Della BellaV, BazzantiM, ChiarottiF. 2005. Macroinvertebrate diversity and conservation status of Mediterranean ponds in Italy: Water permanence and mesohabitat influence. In: Aquatic Conservation: Marine and Freshwater Ecosystems. Vol. 15. pp 583–600.

BiggsJ, WilliamsP, WhitfieldM, NicoletP, WeatherbyA. 2005. 15 Years of pond assessment in Britain: Results and lessons learned from the work of Pond Conservation. In: Aquatic Conservation: Marine and Freshwater Ecosystems. Vol. 15. pp 693–714.

BoyceR. 2009. Fuzzy Set Ordination Web Page. http://www.nku.edu/~boycer/fso/

BrownM, DinsmoreJJ. 1988. Habitat islands and the equilibrium theory of island biogeography: testing some predictions. Oecologia 75:426–9.

Castillo-JuradoM. 1992. Morfometría De Lagos Una Aplicación a Los Lagos.

DowningJ a., PrairieYT, ColeJJ, DuarteCM, TranvikLJ, StrieglRG, McDowellWH, KortelainenP, CaracoNF, MelackJM. 2006. The global abundance and size distribution of lakes, ponds, and impoundments. Limnology and Oceanography 51:2388–97. http://www.aslo.org/lo/toc/vol_51/issue_5/2388.html

EC. 2000. Directive 2000/60/EC of the European Parliament and of the Council of 23 October 2000 establishing a framework for Community action in the field of water policy. Official Journal of the European Parliament L327:1–82.

EdwardsAC, ScalengheR, FreppazM. 2007. Changes in the seasonal snow cover of alpine regions and its effect on soil processes : A review. 163:172–81.

FitterRSR, FitterA, FarrerA. 1984. Grasses, sedges, rushes and ferns of Britain and Northern Europe. London, UK.: Collins

FormanRTT. 1995. Land Mosaics: The Ecology of Landscapes and Regions. 1st ed. Cambridge University Press

García-RollánM. 1985. Claves de la flora de España (peninsula y baleares). Vol.II: dicotiledoneas 20 (l-z) / monocotiledoneas. Mundi-Prensa [in Spanish].

GeerlingGW, RagasAMJ, LeuvenRSEW, Van Den BergJH, BreedveldM, LiefhebberD, SmitsAJM. 2006. Succession and rejuvenation in floodplains along the river Allier (France). In: Hydrobiologia. Vol. 565. pp 71–86.

GoebelPC, PregitzerKS, PalikBJ. 2006. Landscape hierarchies influence riparian ground-flora communities in Wisconsin, USA. Forest Ecology and Management 230:43–54.

GopalB, W.J.J, Davis.JA, editors. 2000. Biodiversity in wetlands: assessment, function and conservation. Leiden, Netherlands: Backhuys Publishers

GottfriedM, PauliH, ReiterK, GrabherrG. 1999. A fine-scaled predictive model for changes in species distribution patterns of high mountain plants induced by climate warming. Diversity and Distributions 5:241–51.

Grey-WilsonC, BlameyM. 1979. The alpine flowers of Britain and Europe. London, UK.: Collins

GwataET, MzezewaJ. 2013. Optional crop technologies at a semi-arid ecotope in southern Africa. Journal of Food, Agriculture and Environment 11:291–5.

HindenH, OertliB, MenetreyN, SagerL, LachavanneJB. 2005. Alpine pond biodiversity: What are the related environmental variables? In: Aquatic Conservation: Marine and Freshwater Ecosystems. Vol. 15. pp 613–24.

HongS-K, KimS, ChoK-H, KimJ-E, KangS, LeeD. 2004. Ecotope mapping for landscape ecological assessment of habitat and ecosystem. Ecological Research 19:131–9. http://www.springerlink.com/index/10.1111/j.1440-1703.2003.00603.x

KamenikC, SchmidtR, KumG, PsennerR. 2001. The Influence of Catchment Characteristics on the Water Chemistry of Mountain Lakes. Arctic, Antarctic, and Alpine Research 33:404–9.

KellerF, GoyetteS, BenistonM. 2005. Sensitivity analysis of snow cover to climate change scenarios and their impact on plant habitats in alpine terrain. Climatic Change 72:299–319.

KesslerJ, ChambraudA. 1990. Météo de la France. Tous les climats, localité par localité. (LattèsJ-C, editor.). Paris, France (in French)

KopacekJ, StuchlikE, StraskrabovaV, PsenakovaP. 2000. Factors governing nutrient status of mountain lakes in the Tatra Mountains. Freshwater Biology 43:369–83.

KörnerC. 1992. Response of alpine vegetation to global climate change. Catena Verlag Supplement:85–96.

KörnerC. 2003. Alpine plant life: functional plant ecology of high mountain ecosystems.

LintingM, MeulmanJJ, GroenenPJF, van der KoojjAJ. 2007. Nonlinear principal components analysis: introduction and application. Psychological methods 12:336–58.

LiuY, PanX. 2014. Ecotope-based Urban Post-industrial Landscape Design. IERI Procedia 9:185–9. http://www.sciencedirect.com/science/article/pii/S2212667814001087

LososJB, RicklefsRE, editors. 2009. The Theory of Island Biogeography Revisited. Princeton University Press

MacArthurRH, WilsonEO. 1963. An Equilibrium Theory of Insular Zoogeography. Evolution 17:373–87.

Mate. 2002. Atlas du Parc Natioal des Pyrénées. GIP ATEN, Morgan Multimedia, EDATER.

MazerolleMJ, DesrochersA, RochefortL. 2005. Landscape characteristics influence pond occupancy by frogs after accounting for detectability. Ecological Applications 15:82434.

Van der MolenDT, GeilenN, BackxJ, JansenBJM, WolfertHP. 2003. Water Ecotope Classification for integrated water management in the Netherlands. European Water Management Online 2003:1–14.

OertliB, JoyeDA, CastellaE, JugeR, CambinD, LachavanneJB. 2002. Does size matter? The relationship between pond area and biodiversity. Biological Conservation 104:59–70.

OksanenJ, BlanchetFG, KindtR, LegendreP,O’HaraRB, SimpsonGL, SolymosP, StevensMHH, WagnerH. 2012. vegan: Community Ecology Package. R package version 1:R package version 2. 0–4.

PaegelowC. 2008. Pyrenäen Bibliografie. Andorra, spanische & französische Pyrenäen. Pyrenees Bibliography. Andorra, Spain & French Pyrenees. Bremen: Verlag Claus Paegelow

Richards-ZawackiCL. 2009. Effects of slope and riparian habitat connectivity on gene flow in an endangered Panamanian frog, Atelopus varius. Diversity and Distributions 15:796806. http://doi.wiley.com/10.1111/j.1472-4642.2009.00582.x. Last accessed 23/07/2011

RichterDD, BillingsSA. 2015. One physical system: Tansley’s ecosystem as Earth’s critical zone, Tansley review. New Phytologist 206:900–12.

RieraJL, MagnusonJJ, KratzTK, WebsterKE. 2000. A geomorphic template for the analysis of lake districts applied to the Northern Highland Lake District, Wisconsin, U.S.A. Freshwater Biology 43:301–18. http://doi.wiley.com/10.1046/j.1365-2427.2000.00567.x

Roberts, D. 2012. labdsv: Ordination and Multivariate Analysis for Ecology. R package version 15-0 http://CRANR-project.org/package=labdsv.

RobertsDW. 1986. Ordination on the basis of fuzzy set theory. Vegetatio 66:123–31.

RobertsDW. 2007. FSO: fuzzy set ordination. R package. http://cran.r-project.orgi.

RobertsDW. 2008. Statistical analysis of multidimensional fuzzy set ordinations. Ecology 89:1246–60.

RobinsonCT, KaweckaB. 2005. Benthic diatoms of an Alpine stream/lake network in Switzerland. Aquatic Sciences 67:492–506. http://www.springerlink.com/index/10.1007/s00027-005-0783-4. Last accessed 30/10/2011

SarosJE, InterlandiSJ, DoyleS, MichelTJ, WilliamsonCE. 2005. Are the Deep Chlorophyll Maxima in Alpine Lakes Primarily Inducedby Nutrient Availability, not UV Avoidance? Arctic, Antarctic, and Alpine Research 37:557–63.

Sorosjinda-NunthawarasilpP, BhumiratanaA. 2014. Ecotope-based entomological surveillance and molecular xenomonitoring of multidrug resistant malaria parasites in anopheles vectors. Interdisciplinary Perspectives on Infectious Diseases 2014.

VernadskyV. 1926. Biosfera. 1st ed. Leningrad, Russia: Nauch

VuilleumierF. 1970. Insular biogeography in contenintal regions. I. The northern Andes of South America. The American Naturalist 104:373–88.

WalkerDA, WalkerDA, HalfpennyJC, HalfpennyJC, WalkerMD, WalkerMD, WessmanCA, WessmanCA. 1993. Long-term studies of snow-vegetation interactions. BioScience 43:287–301.

WhittakerRH, LevinSA, RootRB. 1973. Niche, habitat, and ecotope. The American Naturalist 107:321–38.

WilliamsonCE, SarosJE, SchindlerDW. 2009. Sentinels of Change. Science 323:887–8.

YueTX, LiQQ. 2010. Relationship between species diversity and ecotope diversity. Annals of the New York Academy of Sciences 1195.

ZaharescuDG. 2011. Landscape ecology and geochemistry of high altitude lakes. Insights from the central Pyrenees. PhD thesis, University of Vigo (Spain).

